# The unbiased estimation of *r*^2^ between two sets of noisy neural responses

**DOI:** 10.1101/2021.03.29.437413

**Authors:** Dean A. Pospisil, Wyeth Bair

## Abstract

The Pearson correlation coefficient squared, *r*^2^, is often used in the analysis of neural data to estimate the relationship between neural tuning curves. Yet this metric is biased by trial-to-trial variability: as trial-to-trial variability increases, measured correlation decreases. Major lines of research are confounded by this bias, including the study of invariance of neural tuning across conditions and the similarity of tuning across neurons. To address this, we extend the estimator, 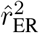, developed for estimating model-to-neuron correlation to the neuron-to-neuron case. We compare the estimator to a prior method developed by Spearman, commonly used in other fields but widely overlooked in neuroscience, and find that our method has less bias. We then apply our estimator to the study of two forms of invariance and demonstrate how it avoids drastic confounds introduced by trial-to-trial variability.

**Significant Statement:** Quantifying the similarity between two sets of averaged neural responses is fundamental to the analysis of neural data. A ubiquitous metric of similarity, the correlation coefficient, is attenuated by trial-to-trial variability that depends on a variety of irrelevant factors. Spearman recognized this problem and proposed corrected methods that have been extended over a century. We show this method has large asymptotic biases and derive a novel estimator to overcome this. Despite the frequent use of the correlation coefficient in neuroscience, consensus on how to address this fundamental statistical issue has not been reached. We both explicate this issue in a neuroscience setting while at the same time making major strides in addressing it.

## Introduction

The measurement of correlation is ubiquitous in sensory neuroscience. The *r*^2^ between two sets of mean neural responses is fundamental to many lines of research including the study of invariance, the maintenance of a tuning curve within a neuron despite the transformation of stimuli (Popovkina et al., 2019; El-Shamayleh and Pasupathy, 2016; Nandy et al., 2013). Transformations can be geometric such as translation (Nandy et al., 2013) or scaling (El-Shamayleh and Pasupathy, 2016) but also can be changes in the ‘appearance’ of a stimuli such as surface luminance (Popovkina et al., 2019). Studies such as these help provide insight into how perception of abstract categories is supported. We show below using the naive estimator, 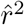, can drastically underestimate invariance. In addition, the correlation between tuning curves has been used for functional clustering providing insights into shared selectivity in and across brain regions (Kiani et al., 2015; Power et al., 2011). Finally, signal correlation, the degree to which tuning curves between neurons correlate, and its relationship to noise correlation and robust encoding has been studied intensively both theoretically and experimentally (Cohen and Kohn, 2011).

In spite of the importance of estimating the correlation across tuning curves, the typical estimator, 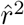, is fundamentally confounded by the trial-to-trial variability of neural responses. Even if two neurons have identical tuning curves, their measured correlation to each other’s responses will decrease as trial-to-trial variability increases. Investigators have approached this problem by averaging over many repeated trials of the same stimulus in order to reduce the influence of trial-to-trial variability, but added data collection is expensive and this approach never wholly removes the influence of noise and its confounding effect. A more principled approach has been to account for trial-to-trial variability in the estimation of correlation. Correction for the attenuation of the correlation coefficients by measurement error has received considerable attention from fields outside of neuroscience (Saccenti et al., 2020; Adolph and Hardin, 2007; Rosner and Willett, 1988; Beaton et al., 1979; Thouless, 1939). The form of the most common correction, given by Spearman (1904), multiplies the correlation coefficient by the inverse of its estimated attenuation. Here we follow an approach developed in the neuroscience literature (Pospisil and Bair, 2020; Haefner and Cumming, 2008; Sahani and Linden, 2003) that removes the bias from the sample covariance and variance separately then forms their ratio for the corrected estimate. We call our estimator 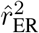 as it estimates the fraction of variance explained between the expected responses 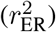 of the neurons or equivalently their tuning curves. We first validate this estimator in simulation. We then compare it to the prior method provided by Spearman and show it has far less bias both for small samples and asymptotically. We then validate our estimator on neural data by showing it, on average, estimates 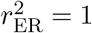 between split trials of the same neuron. Finally we show in neural data how it provides insights into two forms of invariance in area V4. In the first, translation invariance, we find using 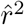 that V4 would appear, on average, to be very sensitive to small shifts in position, but when correcting for noise, with 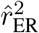, we find selectivity is robust to small shifts. In the second example we study invariance to whether a shape has a uniform surface or is just the outline. We find many neurons are not driven strongly enough to accurately infer invariance, but of those that were driven well by both sets our estimator shows a uniform distribution across the population with some maintaining near identical selectivity while others are wholly orthogonal.

## Materials and Methods

### Simulation procedure

To demonstrate the bias that noise imparts on neuron-to-neuron correlation, and the ability of our methods to remove this bias, we generate values, *r*_*i,j*_, that represent the square-root (for reasons described below) of single-neuron responses to the *j*th repeat of the *i*th stimulus from a normal distribution,

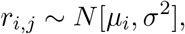

where the mean, *μ*_*i*_, varies across stimuli while the variance, *σ*^2^, is constant. The degree of variation across the *μ*_*i*_ reflects how well the stimuli modulate the neural response, we call this *dynamic range* and quantify it as,

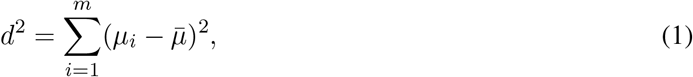

where 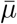 is the average of the *μ*_*i*_ across stimuli. Trial-to-trial variability can be different across cells, fixing *n* and *m*, the signal-to-noise ratio (SNR) determines the reliability with which correlation can be estimated across neuron pairs:

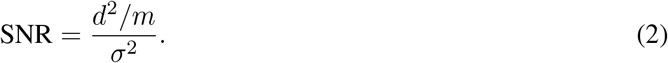

This will be a critical parameter in both our simulations and our analysis of neural recordings.

For our simulations the tuning curves for a pair of neurons, *X* and *Y*, to a set of *m* stimuli are defined by a set of mean responses, *μ*_*i*_ and *ν*_*i*_, respectively, that are modeled as sinusoids (Figure 1, lines), as follows:

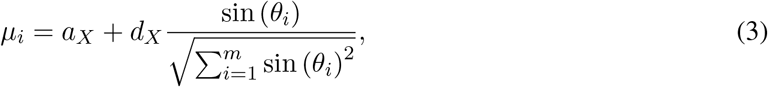

where the denominator normalizes the length of the tuning vector and

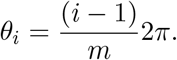

**Figure 1:**
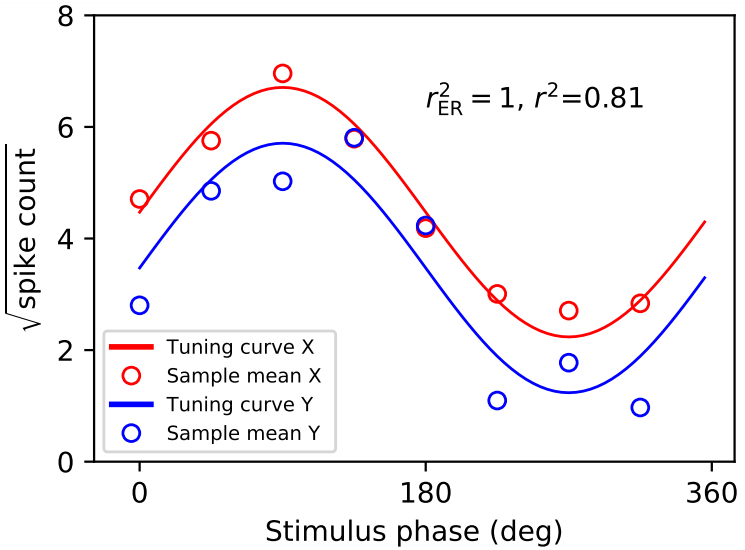
Simulation model of neuron-to-neuron fits. Here 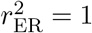 because tuning curves (solid trace) are identical up to a shift and scaling. The estimate of correlation from trial averages (open circles) is lower than 1.

The tuning curve for neuron Y is simply phase shifted to achieve the desired value of 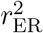, as follows,

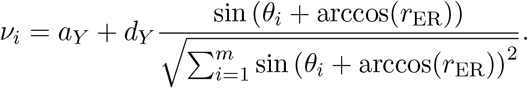

To validate potential estimators for 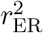, we draw responses independently from this model for *n* repeats for each of the *m* stimuli. Averaging these observations across repeats produces observed tuning curves for the two simulated neurons (Figure 1, open circles).

### Assumptions for the derivation of unbiased estimators

To simplify the derivation of unbiased estimators for 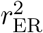, we assume that neural responses have undergone a variance-stabilizing transform such that trial-to-trial variability is the same across all stimuli. For example, if the neural responses are Poisson distributed, *X*_*i,j*_ ~ *P* (*λ*_*i*_), where *X*_*i,j*_ is the response to the *j*th repeat of the *i*th stimulus, which has expected response *λ*_*i*_, then the square root is a variance-stabilizing transform. In particular, if 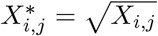, then while the mean still increases with *λ*_*i*_,

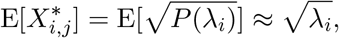

the variance is now approximately constant:

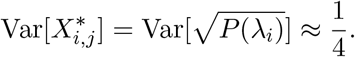

To improve the estimate of the mean response, *n* repeats of each stimulus are collected. Invoking the central limit theorem, we can make the approximation:

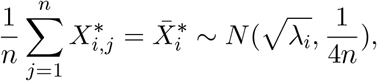

where the average across the *n* repeats is approximately normally distributed with variance decreasing with *n*. The assumption of a Poisson distributed neural response is not always accurate. A more general mean-to-variance relationship,

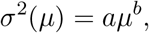

can be approximately stabilized to 1 by,

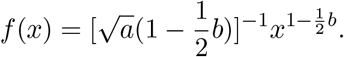

A square root will stabilize any linear mean-to-variance relationship (*b* = 1), but an unknown slope, *a*, requires that this parameter be estimated. In the case of the linear relationship, this simply requires taking a square root and then averaging the estimated variance, which is constant, across all stimuli. If it is not reasonable to assume a parametric mean-to-variance relationship and there are enough repeats, one can simply divide all responses to a given stimulus by their sample standard deviation to achieve *σ*^2^ ≈ 1. For the derivation below, we assume that variance-stabilized responses for each neuron to *n* repeats have been averaged for each of *m* stimuli to yield the mean response to the *i*th stimulus: 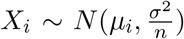 and 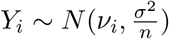, where *σ*^2^ is the trial-to-trial variability and *μ*_*i*_, *ν*_*i*_ the *i*th expected values.

### Unbiasing *r*^2^

In the case where both *X* and *Y* are equal variance stochastic responses, 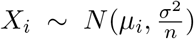 and 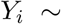 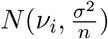, we aim to unbias,

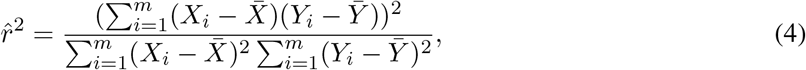

to ideally achieve a corrected estimator, 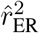, such that,

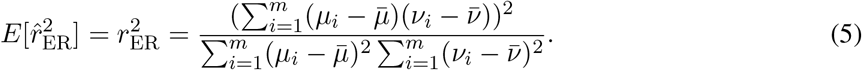

In our approach, we will unbias the numerator and denominator of Eqn. 4 separately. The expected value of the numerator is,

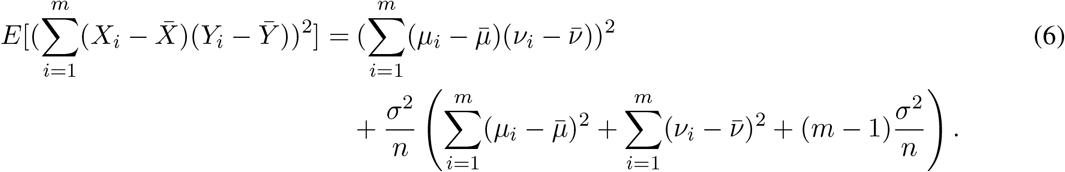

of which the first term is the desired numerator (that of Eqn. 5) and the term on the second line is the bias. To remove this bias an unbiased estimate of it can be subtracted off from the numerator of Eqn. 4.

The denominator of Eqn. 4, consists of two factors that are each scaled non-central chi-squared distributions:

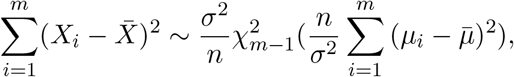

and

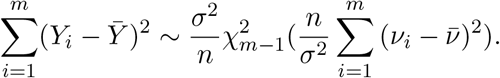

Thus, the expected values of these factors are,

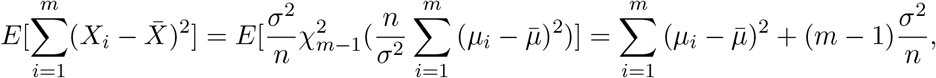

and

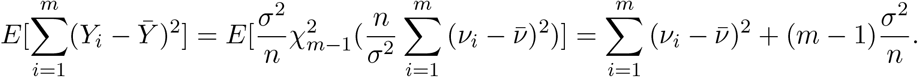

Since these two random variables are independent, the expected value of the denominator, which is their product, is the product of these expected values,

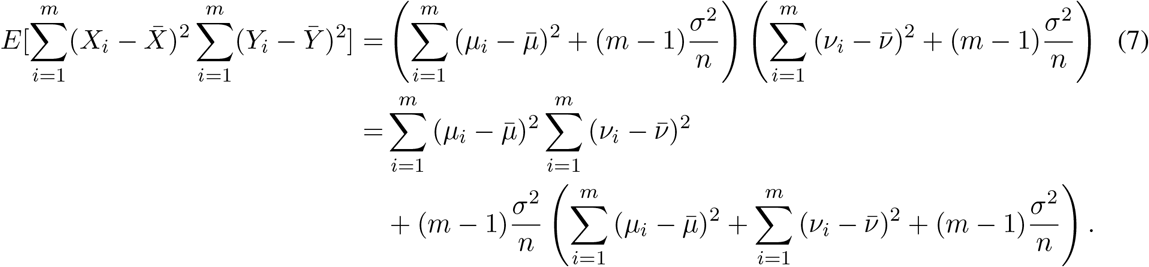

The first term (second line) is the desired denominator (that of Eqn. 5), whereas the second term (third line) is the bias. To remove this bias an unbiased estimate of it can be subtracted off from the denominator of Eqn. 4.

### Estimators of bias terms

To compute the bias terms to be subtracted from the numerator and denominator, three unknown quantities need to be estimated: two dynamic range values,

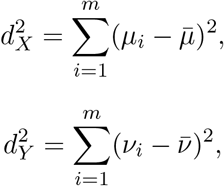

and the trial-to-trial variability, *σ*^2^. Below we provide unbiased estimators for these quantities.

In the case of the dynamic range values, the naive sample estimator, for example,

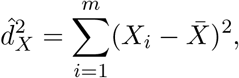

has a bias, as revealed by taking its expected value:

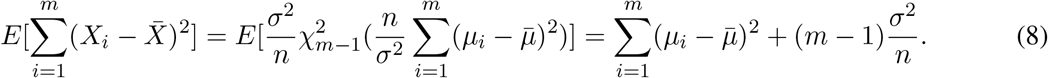

Thus, an unbiased estimator is,

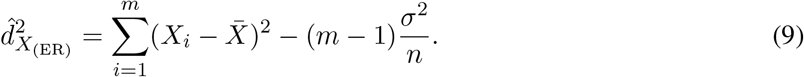

For the case of the sample variance, if we have *n* repeated trials we can calculate sample variance over those trials, then since the variance is the same across stimuli and neurons (assuming a variance stabilizing transformation), we can average them for a global estimate,

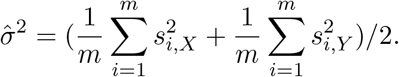

Where 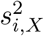 and 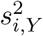 are the sample variance estimates calculated across *n* repeats for the *i*th stimulus. Using these unbiased quantities to form the terms to be subtracted from the numerator and denominator of the naive 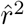 (Eqn. 4), we finally arrive at our corrected estimator for 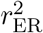, the square of the neuron-to-neuron correlation:

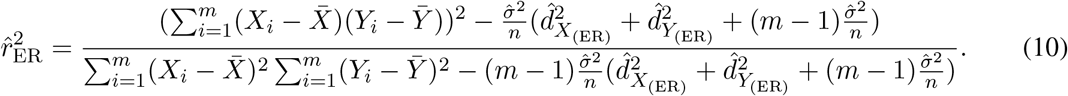

### Spearman’s correction for attenuation

Spearman (1904) noted that in the case of measurements of two quantities with some underlying ‘true’ correlation but with additive independent measurement error, the measured correlation would tend to be less than the true correlation. He provided the analytic expression for the attenuation, *A* (using the notation of Saccenti et al., 2020), under a bivariate normal model,

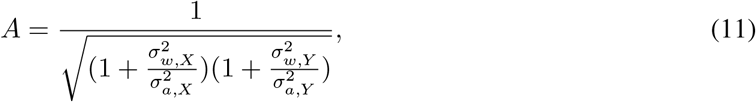

such that the observed correlation, *ρ*, was a scaling of the true correlation, *ρ*_0_,

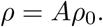

Here, 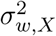 and 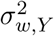 are the variance of the additive measurement error (within-condition variance) and 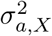 and 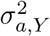 are the variance of the underlying quantities (across-condition variance). This is slightly different than our derivation above. Here, both the correlated and noisy components are random, whereas in the previous section our correlated component was fixed and only the noise component was random. Thus we use 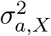 instead of 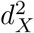, because the former is the variance of a normal distribution whereas the latter is the squared deviation of a set of means

The method to reverse attenuation is straightforward: find an estimate for *A* and multiply estimated correlation, *ρ*_0_, by the inverse of this estimate. Spearman does not specify estimators for the unknowns in Eqn. 11, however, Adolph and Hardin (2007) use sample variance to estimate both the within-condition variance (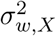 and 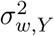) and across condition-variance (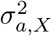 and 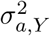). Thus, using the estimators defined in the previous section and assuming equal noise variance 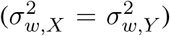, Spearman’s corrected estimator takes the form,

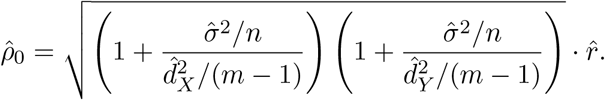

We square this estimator to compare it to ours. Because 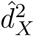 and 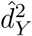 are biased estimators (see Eqn. 8), we also use the unbiased estimators shown in Eqn. 9. We call this estimator 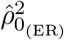.

### Electrophysiological data

To demonstrate the use of our estimator, we re-analyzed data from three previous single-unit, extracellular studies of parafoveal V4 neurons in the awake, fixating rhesus monkey (Macaca mulattta). Data from the first study, (Pasupathy and Connor, 2001), consists of the responses of 109 V4 neurons to a set of 362 shapes. There were typically 3-5 repeats of each stimulus, and we used only the 96 cells that had at least 4 repeats for all stimuli. We used the spike count for each trial during the 500 ms stimulus presentation. From a second study, (El-Shamayleh and Pasupathy, 2016), we re-analyzed responses of 80 neurons tested for translation invariance using shapes like those of the first study, but where the position of the stimuli within the receptive field (RF) was also varied. Each neuron was tested with up to 56 shapes (some of which are rotations of others) presented at 3-5 positions within the RF. Each unique combination of stimulus and RF position was presented for 5-16 repeats, and spike counts were averaged over the 300 ms stimulus presentation. Data from the third study, (Popovkina et al., 2019), consists of the responses of 42 V4 neurons using shapes like those of the first study except in two conditions: fill and outline. In the fill condition, the interior of the shape was the same color as its outline. In the outline condition, the interior of the shape was the same color as the background. Stimuli were presented for 300 ms and a 200 ms blank interval preceding each. For each neuron, stimuli were presented for at least 3 repeats.

Experimental protocols for all studies are described in detail in the original publications.

### Software Accessibility

Code for calculating estimates of 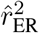 and associated confidence intervals are available at https://github.com/deanpospisil/er_est.

## Results

Our results are organized as follows. We first describe the source of the bias in 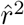 using idealized neuronal tuning curves and show how our corrected estimator, 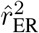, removes this bias. We next validate our method in simulation, comparing it to alternative methods, and we validate it using neural data in a split-trial comparison. Finally, we demonstrate how our estimator avoids significant confounds in two applications: the measurement of translation invariance and fill-outline invariance in single-unit data collected in previous studies of shape selectivity in visual cortical area V4.

Consider a typical scenario in sensory neuroscience where the response of two neurons to *n* repeated presentations of *m* stimuli have been collected and the average of these responses are compared (Figure 1). Even if the underlying neuronal tuning curves (blue and red lines) were perfectly correlated, the *m* sample averages, *Y*_*i*_ and *X*_*i*_ (blue and red open circles), will deviate from the expected value owing to trial-to-trial variability, *σ*^2^, scaled by 1*/n*. Thus, the observed 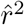 can be appreciably less than 1 even though the *r*^2^ between the *expected values* of the neuronal responses is 1 (red and blue lines are identical up to a shift and scaling).

The quantity we attempt to estimate here is the correlation coefficient *r*^2^ between the expected values, *μ*_*i*_ and *ν*_*i*_, of the responses of the two neurons, i.e., between the tuning curves in the absence of noise. We will call this quantity 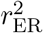, the fraction of variance shared by the expected responses (ER) of the two neurons,

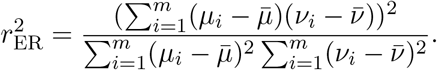

It is tempting to estimate this quantity using the naive sample estimator,

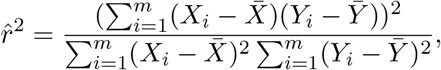

however, the expected values of the numerator and denominator of 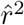 have a bias with respect to those of 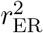, as shown in Methods Eqns. 6 and 7, respectively. Each has a bias proportional to *σ*^2^*/n*, the amount of variability remaining after averaging over stimulus repeats. The bias in the denominator, which results in the attenuation of 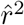, remains even as *m* → ∞ because the effect of trial-to-trial variability is accumulated across stimuli in the calculation of total variance in the denominator. This is reflected in the scaling of the denominator’s bias by (*m* − 1) (Eqn. 7, third line). The bias does go to zero as *n* → ∞ because averaging across repeated presentations of stimuli reduces the effect of trial-to-trial variability. This is reflected by the trial-to-trial variability factors in the bias of both Eqns. 6 and 7 being scaled by 1*/n*.

To solve this problem, we take the straightforward strategy of finding unbiased estimators of these noise terms and subtracting them from the numerator and denominator of 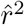 to give our estimator 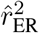 (see Methods, Eqn. 10).

### Validation of estimator by simulation

To test the ability of 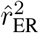 to accurately estimate true neuron-to-neuron tuning curve correlation in the face of noise, we simulated the responses of two neurons with the same trial-to-trial variability (*σ*^2^), the same number of trials (*n*), but potentially different SNR (Eqn. 2). Both neurons had sinusoidal tuning curves (as in Figure 1), and their relative phase offset was varied to set the true tuning correlation (Methods, Eqn. 3). We first consider the case where neuron *X* has a high SNR (SNR_X_ = 1.0) while neuron *Y* has a low SNR (SNR_Y_ = 0.5). In this case, the naive estimate of correlation, 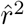 (Figure 2A, orange), is on average only one quarter of the value of the true correlation between the simulated tuning curves. For example, 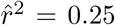 when 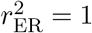. On the other hand, our 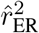 on average lies very close to the true correlation (Figure 2A, blue line lies on diagonal). However, although 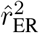 is clearly the less biased estimator of 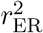, it has become more variable: the 95% quantile error bars are longer than those for the naive estimator.

**Figure 2:**
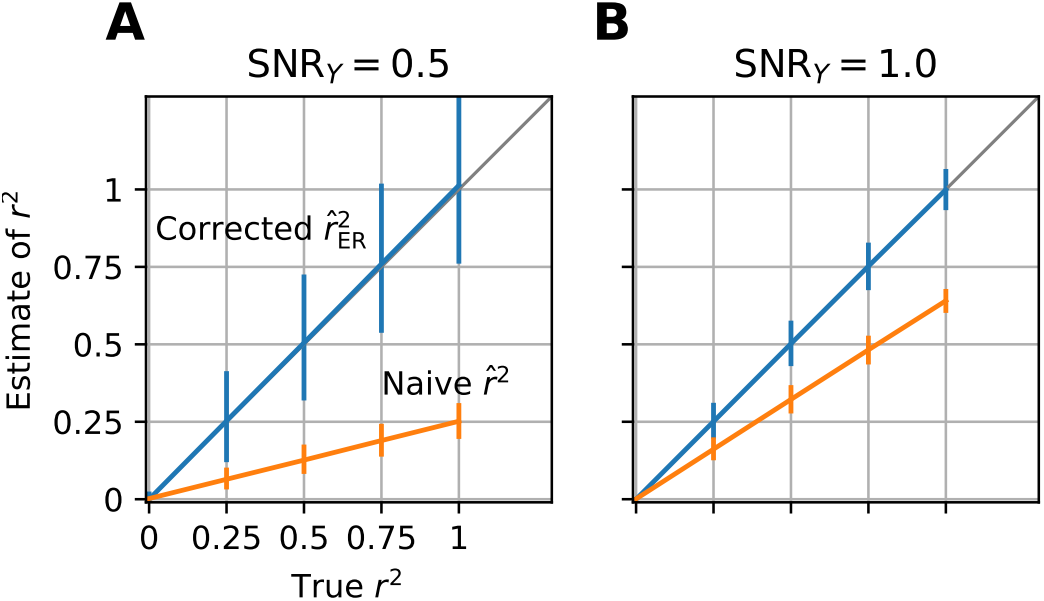
Simulation of 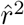 (orange) and 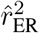 (blue) for estimating fit of neuron-to-neuron at varying levels of *r*^2^ and SNR of neuron Y, SNR_Y_, while the other neurons SNR_X_ = 1 stays fixed. Number of trials are *n* = 4, stimuli *m* = 371, and trial-to-trial variability is set at *σ*^2^ = 0.25. Vertical bars are 95 % quantiles. **(A)** Simulation at lower SNR, SNR_Y_ = 0.5 and SNR_X_ = 1. **(B)** The same simulation as (A) but both neurons have SNR=1.

When both neurons have a high SNR (Figure 2B) the difference between the average of 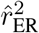 and 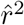 shrinks commensurate with the increased SNR, as does the difference in the error bars.

### Comparison to Spearman’s correction

Here we compare our estimator, 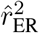, to that of Spearman as formulated by Adolph and Hardin (2007), 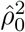, and an extension, 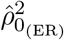, where we substituted unbiased estimates of 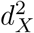 and 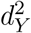 (see Methods, ‘Spearman’s correction for attenuation’). For reference, we also compare these estimator to the naive estimate 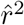.

We compare these four estimators using simulated data where we vary the SNR (Eqn. 2), equal for both neurons, and the number of stimuli *m* (Figure 3). In the case of a small number of stimuli (*m* = 20), the naive 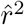 is well below 1 and remains below for all levels of SNR (orange trace Figure 3A). This bias remains even as *m* is increased to 50 (panel B, orange) and 1000 (panel C, orange). The estimator proposed by Adolph and Hardin, 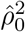, is less biased (red trace) but still consistently underestimates the true *r*^2^ and does not improve with *m*. The same estimator computed using an unbiased estimate of the dynamic range, 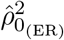, shows an upward but smaller bias relative to the downward bias of 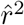 and 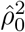. It converges to the true value with *m* (Figure 3, purple). Finally, 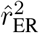 (blue trace) shows the least bias of the four estimators over a wide range of SNR and *m*, though in the case of the lowest SNR values it becomes unstable (panel A, large blue error bars). For higher values of *m* (panels B and C) this is not an issue. For very high *m*, our correction to Spearman’s method 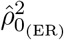 and our estimator 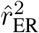 are essentially identical (blue and purple lines overlap, Figure 3C).

**Figure 3:**
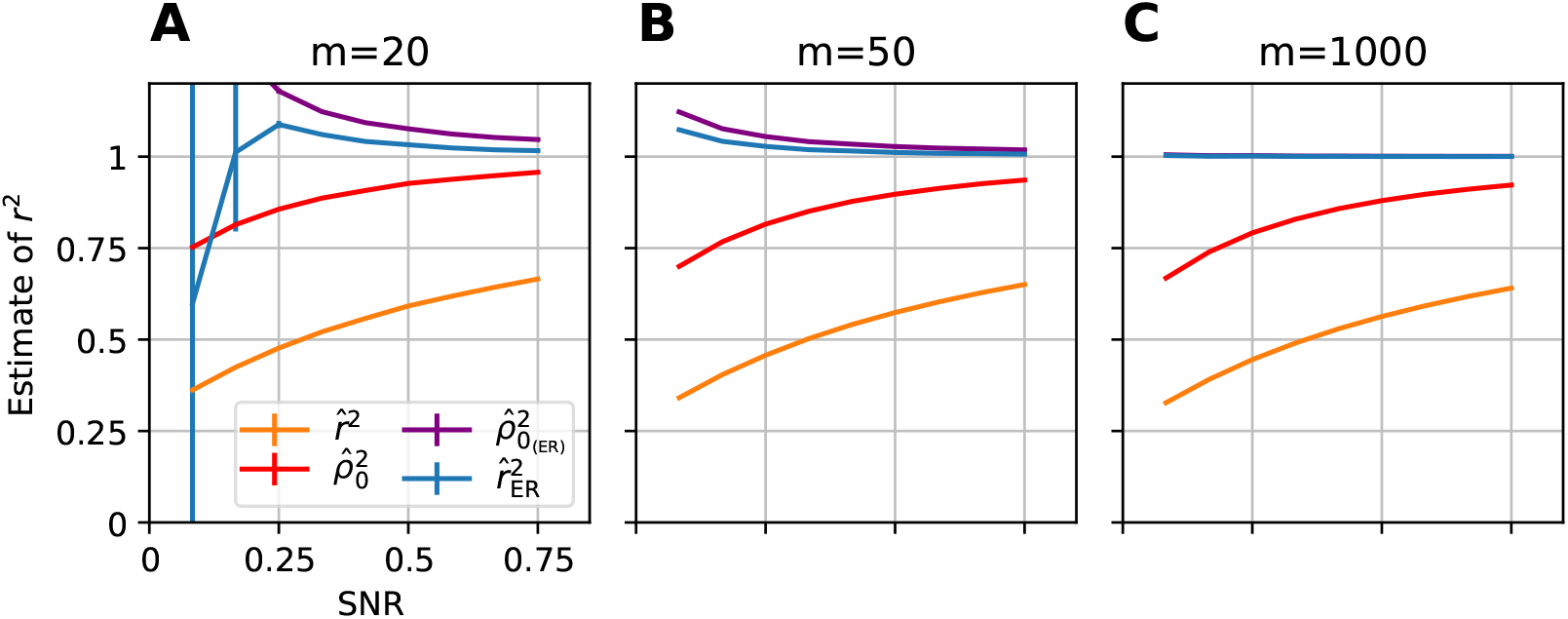
Comparison of 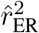 to alternative methods. The mean and SE (vertical bar) of each estimator is calculated across a simulation of fitting two neurons with *n* = 4 repeats (see Methods, ‘Simulation procedure’) for 10,000 simulations. In orange is the naive estimator Pearson’s *r*^2^, in red is Spearman’s estimator (1904) calculated according to methods of Adolph and Hardin 2007, in purple is Spearman’s estimator with an unbiased estimator of dynamic range Eqn. 9, and in blue is the estimator we use through out the paper, 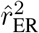. **(A)** Low number of stimuli (*m*). **(B)** Intermediate number stimuli. **(C)** High number of stimuli.

### Split trial validation

We have shown that our estimator, 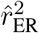, works well under simulations that follow the assumptions of its derivation. Because neuronal responses are not guaranteed to be well approximated by these assumptions, it is important to test the method on real neuronal data. In most cases, we do not know 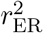 so we cannot determine whether the estimator is working. One case in which we do know 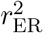 is across independent trials of a neuron’s responses to the same stimuli. Theoretically, the expected values are the same for two tuning curves computed from alternating stimulus repeats recorded from a single neuron, thus 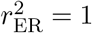. Here we correlate the mean responses from the odd trials to those from the even trials for a set of single neurons with the hope that our method correctly estimates 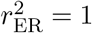. Our neuronal data set consists of single-unit recordings from 96 V4 neurons with 4 repeats (see Methods, ‘Electrophysiological data’). We evaluate our estimators on the 1st and 3rd trial correlated to the 2nd and 4th trial.

We first computed the naive 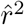 for all cells (Figure 4, orange points) and plotted these values as a function of the estimated SNR (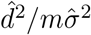, across all trials). Despite the theoretical value of 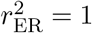, the unit with the highest SNR (rightmost orange point) achieves only 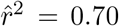. Thus, for real neural data with low *n*, the naive 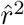 clearly underestimates 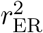. In essence, 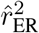 was constructed to correct all of these points to be distributed about 1. Indeed, the scatter of 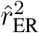 values across the population of cells is approximately centered around 1 (Figure 4, blue points), with an increasing upward bias for low SNR. The highest SNR units (rightmost blue points) have estimates that are close to 1. For neurons with lower SNR, some of the 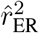 values become unstable, taking on values far greater than 2 (Figure 4, small blue points, top left) or less then 0 (small blue points, bottom left) consistent with our simulations (Figure 3A, blue error bars on left).

**Figure 4:**
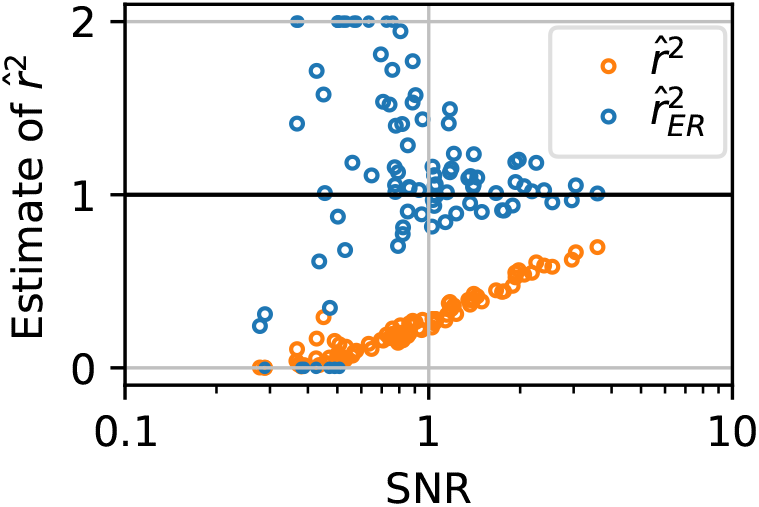
Motivation and validation of 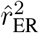 on split-half correlation for neural responses of Pasupathy and Connor (2001). Estimated correlation between the odd and even trials of a neurons responses to a set of *m* =371 stimuli conditions, 96 neurons total. Plotted as a function of the SNR of the neurons calculated across all trials. In blue are the corrected 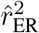 and in orange the naive 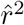. Theoretically 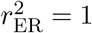 (bold black horizontal line). Corrected estimates that went above 2 are set to 2 or below 0 are set to 0.

For two example neurons, plots of the raw tuning curve values on even vs odd trials (Figure 5) provide deeper intuition into the metric. Each point in the scatter plots represents the mean response to a particular shape on the even trials vs that on the odd trials. In both plots, by the split-half construction, all of the residual variance is attributable to trial-to-trial variability. The estimator 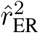, being approximately 1.0 in both cases, is appropriately factoring out the trial-to-trial variability and is in essence predicting that with more trials, one should expect these points to settle onto the diagonal.

**Figure 5:**
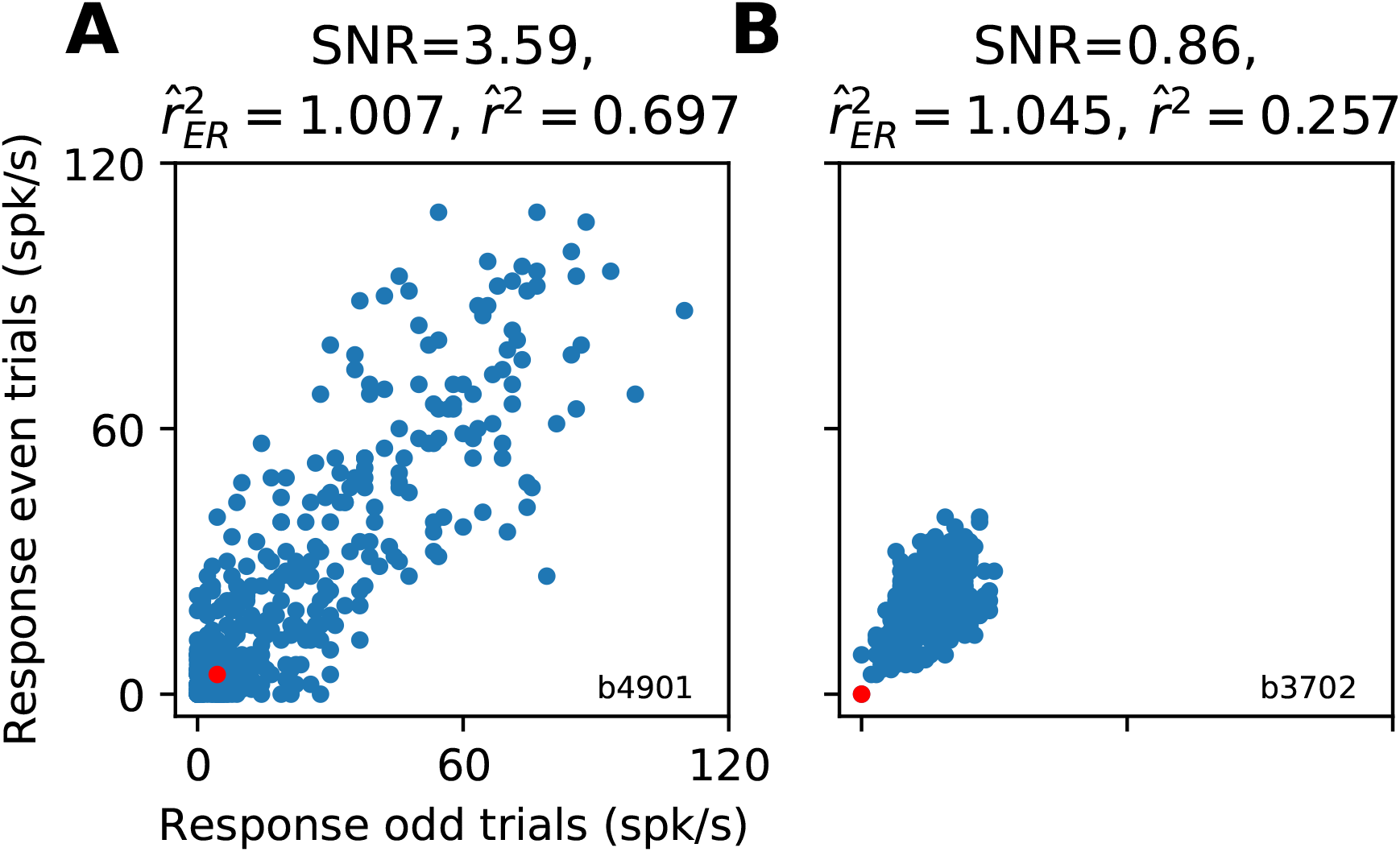
Example cells from split-trial correlation of Pasupathy and Connor (2001). **(A)** Example cell with high SNR. Each point is the average of the even (x-axis) and odd (y-axis) spike rate for a single stimulus. In red is the baseline firing rate. SNR is high because of a large dynamic range in spike count. **(B)** Example cell with smaller SNR. Despite 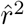 being lower in this case the estimator predicts that, similarly to (A), the neural responses have the same pattern of means.

Overall, our validation on simulated data demonstrates that our corrected 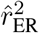 avoids the bias of the naive estimator, 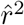, and outperforms a previously proposed metric. Furthermore, our metric appears able to achieve an approximately veridical average result for a population of real neuronal data in a split-half test that had very low *n* (only two repeats) and thus substantial noise. Next, we apply our method to estimate relationships in neuronal tuning where no ground truth exists.

### Measuring population translation invariance

Translation invariance of neuronal selectivity is the degree to which a neuron maintains the same pattern of responses to a stimulus set regardless of where the stimulus set is presented in its RF. This can be quantified by measuring the correlation between the responses at one position to those at another. The naive 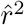 will suffer from the confounds described above. To avoid this, we use 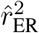 to accurately assess translation invariance in V4. A population level estimate of invariance found by averaging across single neuron estimates will inherit the unbiased quality of the individual samples and thus will provide an accurate estimate of typical translation invariance in V4.

We reanalyzed data from El-Shamayleh and Pasupathy (2016) that consisted of recordings of 80 V4 neurons responding to a set of simple shapes presented at up to 4 positions within the RF (see Methods, ‘Electropysiological data’). One potential concern is that the RFs of the neurons are diverse (Figure 6A, grey traces): for some neurons the average response (across shapes) falls off quickly over space whereas for others the mean remains high. On average, the neurons are at less than ~80% of mean response for the furthest stimuli. Given our study of SNR, it is quite possible that the naive 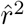 would display a fall off in correlation simply because SNR was lower further from the center of the RF (as in Figure 3, orange trace). Indeed 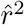 drops off sharply from the center of the RF (Figure 6B), falling from a value of 1 (by definition) at the center to a mean of 0.51 to the left of center and 0.48 to the right of center. The mean 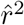 value continues to drop further to the edges of the range tested (0.23 and 0.27 at left and rightmost points, respectively). Knowing that this naive metric is biased downward by noise and depends on SNR raises the distinct possibility that such a sharp drop would not be observed if more trials were collected. To overcome this confound, we used our corrected metric 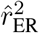 (Figure 6C) and averaged across neurons by weighting each value by the SNR of the associated responses. This reveals that for the smallest shifts away from the center, tuning remains quite similar on average (e.g., 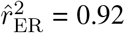 right of center). Thus, the initial steep drop off (in Panel B) is attributable to trial-to-trial variability. On the other hand, a substantial drop off in tuning correlation remains for the largest offsets tested.

**Figure 6:**
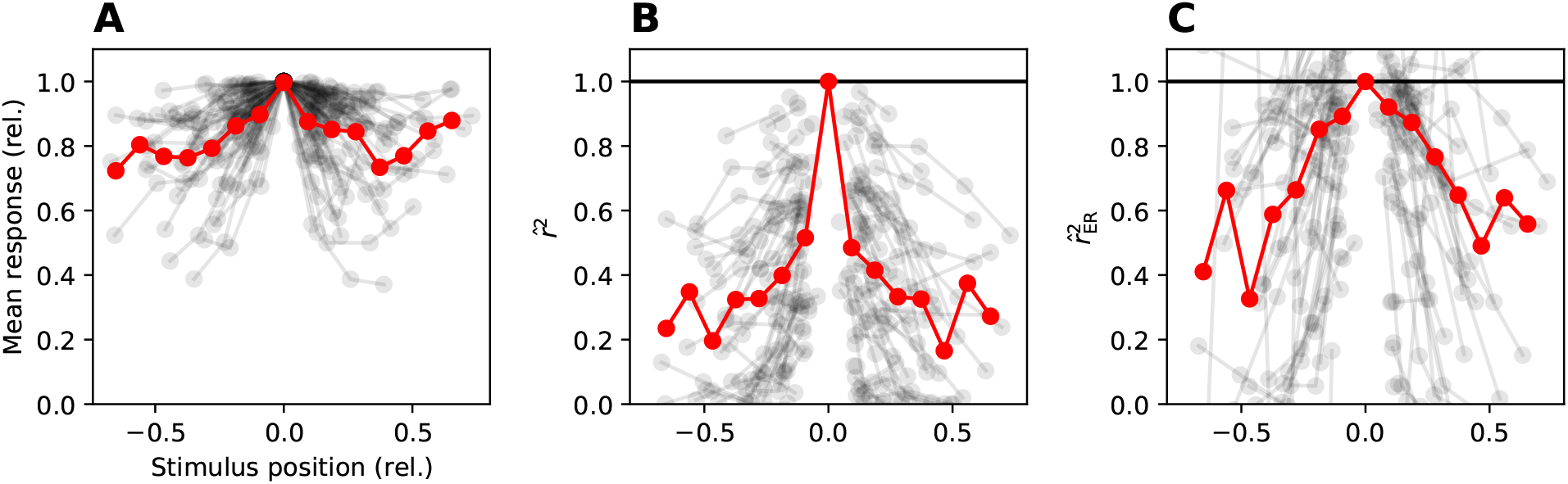
RFs and translation invariance across 80 cell from El-Shamayleh and Pasupathy (2016). **(A)** Average response across stimuli of each cell at each position of stimuli in RF (grey lines) normalized to peak average response defined to be center of RF. Plotted as function of shift of stimuli in units of fraction of RF. Average across population (red) points are the average across the population in 15 non-overlapping bins of 1*/*10 RF width, points are at center of bins. **(B)** The 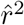 of all cells responses to stimuli at center RF with those at a shift (grey). Center is by definition 1. Average across population with same binning as (*B*) in red. **(C)** 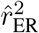 of all cells (grey) and weighted binned average (red, weighted by SNR).

By focusing on neurons with high SNR, we can find units that are truly invariant and we can also identify units that have tuning that changes reliably with position in the RF. For example, Figure 7 shows data for one high-SNR neuron (thus strongly tuned for shapes; neuron 68) with high translation invariance (solid blue trace is relatively flat) and for another (neuron 25) with tuning that is more sensitive to position (solid cyan trace). For the same fraction estimated RF shift (−0.12), one cell has near-perfect invariance, despite the RF dropping off by ~ 11% (blue dotted) whereas the second example cell shares only half the variance with the tuning at the center of the RF (cyan trace; RF drops ~ 20%). The important contribution of the 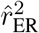 estimator is that it removes ambiguity about whether such differences are caused by changes in firing rate over position. Examining the responses at positions 0 and −0.12 across shapes (Figure 7B, each point corresponds to one shape), for neuron 68 (blue) there is a strong linear relationship, with residuals attributable to trial-to-trial variability. For neuron 25 (cyan), however, there is a substantial change in tuning: many shapes with points previously in the cluster near the origin have more than doubled their responses at the shifted position, whereas for points to the right of 20 spikes/s, the average response to these shapes has been roughly halved.

**Figure 7:**
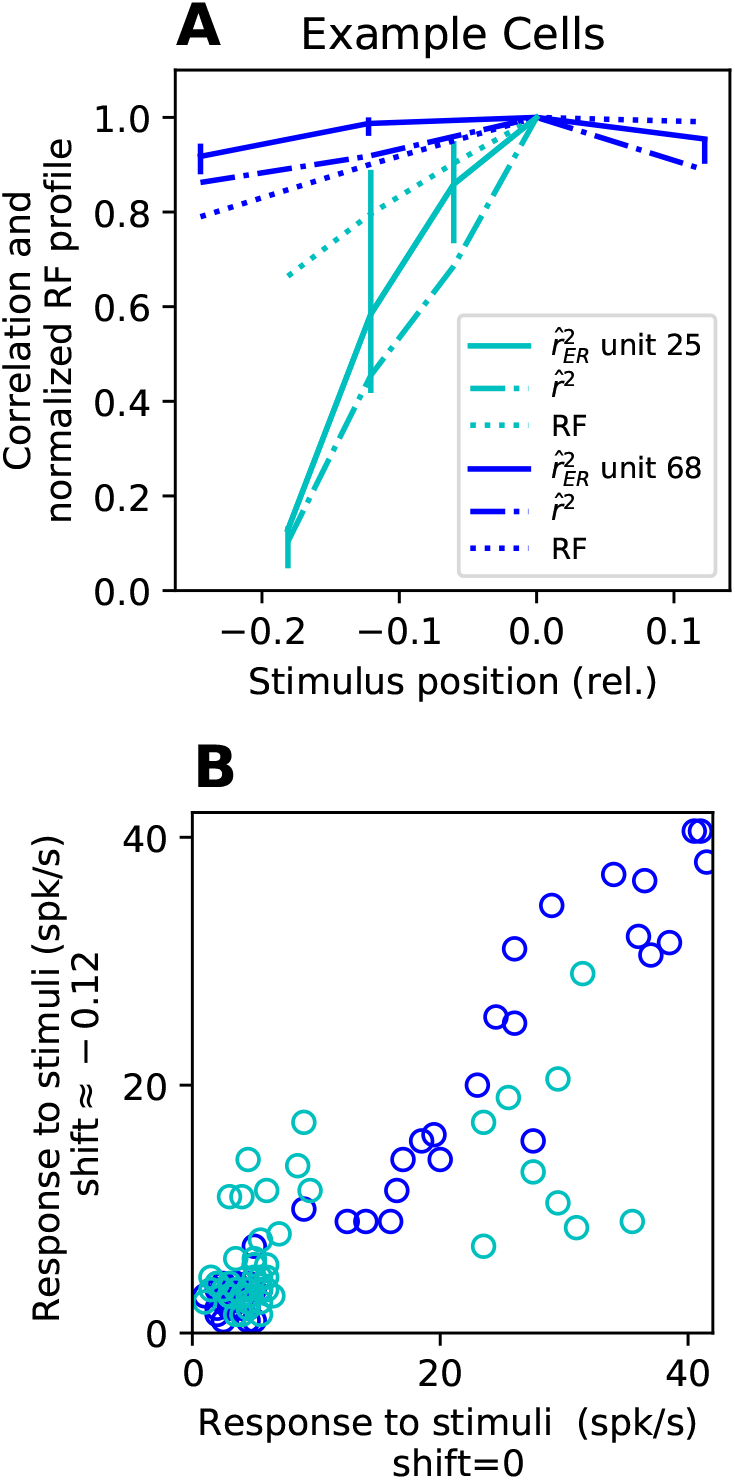
Example neurons showing diversity in translation invariance (TI). Examples were chosen for low and high TI, but high SNR with measurements at similar position in RF. **(A)** A cell with high TI (blue) maintains a high 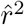 (dash and dot line) across RF (dotted line) and 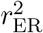 is near 1 reflecting near perfect invariance as would be expected for a high 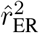. For a second example cell (cyan) 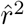 drops off quickly and 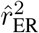 is similar thus the drop is not the result noisy responses. **(B)** Comparing the responses of the two example cells where the shift at −0.12 fraction of RF (2nd point from left on solid lines in (A)) is plotted against the responses at the center. Unit 68 (blue) shows a strong linear relationship reflecting its high 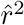 whereas for unit 25 one subset of stimuli evoke a higher response at the shifted position (higher cyan points on left) then at the center and another subset (cyan lower right) evoke a lower response thus tuning is clearly changing with position.

In summary, 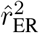 can distinguish with confidence between units that are sensitive vs. insensitive to stimulus position and can provide unbiased averages across a population without needing to resort to removing all but the most well-tuned neurons.

### Measuring correlation between tuning for shapes and their outlines

We now demonstrate how our unbiased metric along with its associated confidence intervals (CIs) can be used to accurately and systematically quantify another important visual invariance, one that relates to the representation of boundary versus surface features. The neuronal data comes from a study (Popovkina et al., 2019) in which V4 neurons were tested with a set of shapes presented as either outlines (shape interior matched the background) or fills (interior was painted the same as the boundary) with the goal of determining whether responses were dictated by boundary shape alone or were dependent on interior fill. It was hypothesized that neuronal responses would have a high correlation across the transformation from fill to outline if neuronal shape tuning was largely based on boundary information, but it turned out that the r-values tended to be low, suggesting that most neurons did not show strong fill-outline invariance. This is an intriguing observation because many models differ strongly from V4 in this respect: for example, an H-Max model fit to V4 shape tuning was found to have very high FO invariance (Popovkina et al., 2019). But, to be able to accurately compare noisy neuronal data to noise-free models, we need a way to correct for noise. To address this problem by applying our 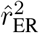 estimator, it is important to consider that the added variability in the corrected estimate can be confounding, as it will tend to smear out the population distribution. To this end, we can make use of confidence intervals to select only those cells with reliable 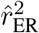 values. For this subset, we can be confident that the empirical distribution is accurate and not degraded by noise. This criterion naturally removes neurons that were not significantly tuned for both sets of stimuli; such neurons that lack measurable tuning for only one condition can be considered as a special case of non-invariance.

We computed fill-outline 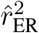 values and associated confidence intervals for 42 V4 neurons and found that nearly 40% of the cells had CIs that spanned the full [0, 1] range (Figure 8A, rightmost bar). These neurons correspond to the observation in the original study that many neurons were not tuned for either the fill or the outline stimulus sets, and this lack of tuning means that the SNR is too low to allow for accurate unbiasing of the correlation value.

**Figure 8:**
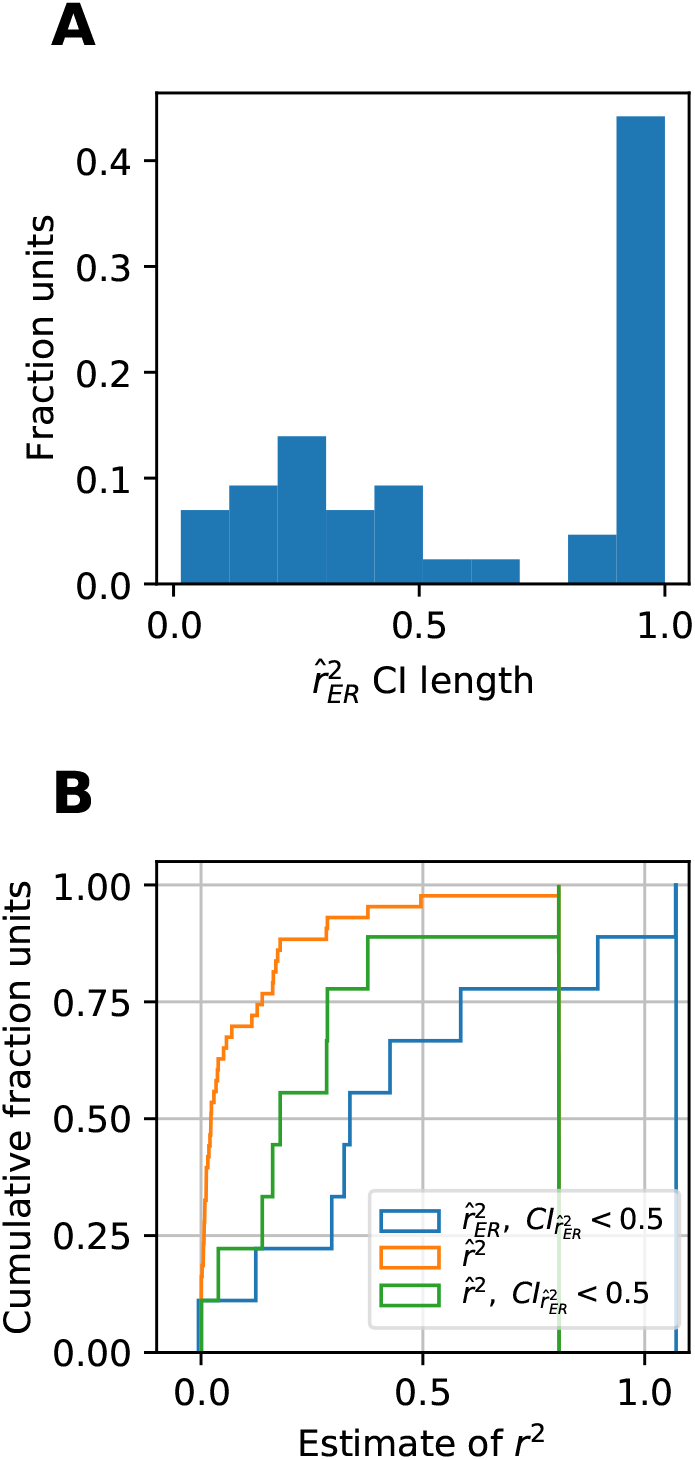
Comparison of 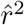 and 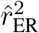 for determining strength of correlation between fill and outline responses. **(A)** The distribution of length of 90 % confidence intervals. **(B)** Cumulative distribution of 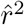 (orange), 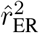 (blue) where only 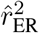 with an associated confidence interval with length less than 0.5 are included leaving 17 cells of the original 42, and the 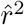 of the same 17 cells (green).

The cumulative distribution of the naive 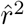 across all neurons (Figure 8B, orange trace) climbs quickly along the left side of the plot, suggesting that high fill-outline invariance is rare (median 0.02). In comparison, the distribution of 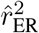 values for those neurons with narrow confidence intervals (blue line, CI*<*0.5) is shifted substantially rightward and becomes broader (median 0.38). In this distribution, neurons that reliably change their pattern of selectivity across the fill and outline conditions are indicated by low values, whereas neurons for which selectivity for either fill or outline stimuli is absent or unmeasurable are now excluded. To allow a direct comparison between the naive and corrected correlation values for the same set of units, the orange line shows the distribution of the naive estimator for just those units with narrow CI. From this comparison, we conclude that units with measurable tuning for both fill and outline stimuli tended to sit on the higher end of the original naive distribution (green curve lies to the right of orange curve), and that our corrected values are substantially larger than the naive values (blue lies to the right of green).

Example neurons can provide some intuition into the corrected values. Figure 9A plots outline vs. fill responses for a cell with middling 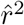 but high 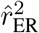 (values shown above panel). The latter value indicates that the observed variation in the scatter plot is consistent with what would be expected given the trial-to-trial noise for this neuron, assuming it had identical fill and outline tuning. Cells with relatively low 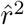 and middling 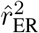 (Figure 9B and C) are those where some of the variance, but not nearly all, is attributable to noise. Finally, a cell with low SNR (Figure 9D) had a confidence interval covering [0,1] consistent with the observation that it responded significantly only to filled shapes and not to outlines.

**Figure 9:**
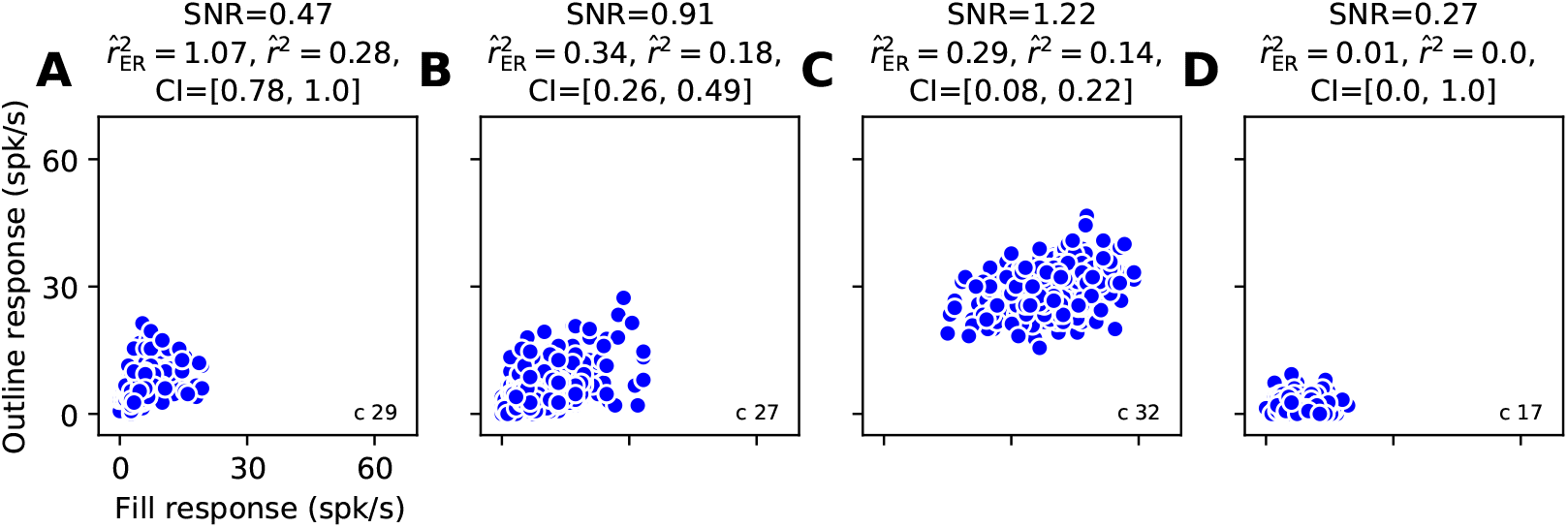
Example cells responses to fill and outline shape stimuli. **(A)** Cell with lower SNR but high fill-outline invariance. On the basis of 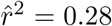 it might be assumed this has low invariance but given the amount of noise and low dynamic range this correlation is quite high and confidence intervals suggest the true invariance is upwards of 0.7. **(B)** Example of cell with high SNR but middling fill-outline invariance. **(C)** Example of cell with higher SNR but little fill-outline invariance. **(D)** Example of cell with low SNR and confidence interval of length 1. Typically of units with long CI’s the neuron only evokes modulation for one set of stimuli (in this case filled shapes).

In summary, by focusing on 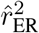 estimates with small confidence intervals, we were able to show that the population of neurons with significant tuning for both fill and outline stimuli displays a broad spectrum of fill-outline invariance and has a higher average invariance then would be predicted based on a naive analysis of all neurons.

## Discussion

### Summary

We have introduced a new estimator, 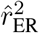, for the fraction of variance shared between the expected value of two neural tuning curves. We show that this estimator has less bias than previous methods of accounting for the attenuation of correlation by trial-to-trial noise. We have demonstrated in two different neural data sets that it avoids ambiguity and confounds that can meaningfully change conclusions that are drawn from the data, particularly in comparison to the commonly-used, naive 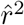. In the analysis of tuning invariance to stimulus transformations, we found 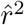 typically grossly underestimated invariance but 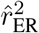 did not. For translation invariance, we showed how 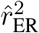 could be used for estimating average population invariance that will be useful for making noise-corrected comparisons across data sets and for comparison to invariance measured in noise-free models. To our knowledge this is the only estimate of population translation invariance in V4 corrected for noise. For fill-outline data, we showed how the confidence intervals could be used as a criterion for including only data points where unbiasing could be reliably achieved. Our metric with associated confidence intervals provides a novel tool for future studies that seek to avoid confounds of noise when comparing responses across stimulus conditions and across neurons.

### Quantifying invariance

We quantified invariance as the correlation in the tuning curves of a neuron to a set of reference stimuli and a set of transformed stimuli. In our case, the transformations were image translation and the removal of all but the outline of a shape. This correlation measure is explicitly insensitive to changes in the mean or amplitude of tuning curves. We consider this a ‘weak’ form of invariance since the spike count across the transformation for a given stimulus can be wildly different. ‘Strong’ invariance would be if the tuning curve does not change at all across a transformation. One condition in which weak invariance may become strong is if gain control mechanism can remove the difference between the two sets of interleaved stimuli when they are shown in separate blocks (Ohzawa et al., 1985). This is a testable hypothesis: do neurons with weak invariance for interleaved stimuli achieve strong invariance when those stimuli are blocked. The estimator we have introduced would be crucial to answering this question.

### Further work

The correlation coefficient is the basis of several multivariate data analysis methods. Examples include canonical correlation analysis (CCA), representational similarity analysis (RSA), and principal component analysis (PCA) when the variance is factored out. With respect to neuroscience, the latter is the basis of a popular measure of neural dimensionality and the method of spike-triggered covariance. Further work can study the effect of noise (specifically in shrinking correlation) on the estimation of these quantities and if applying our corrected estimator can improve inference.

Here our derivations have assumed the case of independent samples, but often nearby neurons are recorded simultaneously and show correlation across trials, termed noise correlation (Cohen and Kohn, 2011). In the future, our estimator could be extended to account for the effect of correlated samples.

## Acknowledgements

We thank Dina Popovkina, Yasmine El-Shamayleh, and Anitha Pasupathy for sharing data. We thank Greg Horwitz and Yen-Chi Chen for providing helpful comments. This work was funded by a National Science Foundation (NSF) Graduate Research Fellowship DGE-1256082 (D.A.P.), NIH/NEI R01-EY02999, and NIH/NEI R01-EY027023 (W.B.).

## References

Pospisil, D. A., & Bair, W. (2020). The unbiased estimation of the fraction of variance explained by a model [Publisher: Cold Spring Harbor Laboratory Section: New Results]. bioRxiv, 2020.10.30.361253. https://doi.org/10.1101/2020.10.30.361253

Saccenti, E., Hendriks, M. H. W. B., & Smilde, A. K. (2020). Corruption of the Pearson correlation coefficient by measurement error and its estimation, bias, and correction under different error models [Number: 1 Publisher: Nature Publishing Group]. Scientific Reports, 10(1), 438. https://doi.org/10.1038/s41598-019-57247-4

Popovkina, D. V., Bair, W., & Pasupathy, A. (2019). Modeling diverse responses to filled and outline shapes in macaque V4. Journal of Neurophysiology, 121(3), 1059–1077. https://doi.org/10.1152/jn.00456.2018

El-Shamayleh, Y., & Pasupathy, A. (2016). Contour Curvature As an Invariant Code for Objects in Visual Area V4. Journal of Neuroscience, 36(20), 5532–5543. https://doi.org/10.1523/JNEUROSCI.4139-15.2016

Kiani, R., Cueva, C. J., Reppas, J. B., Peixoto, D., Ryu, S. I., & Newsome, W. T. (2015). Natural grouping of neural responses reveals spatially segregated clusters in prearcuate cortex. Neuron, 85(6), 1359–1373. https://doi.org/10.1016/j.neuron.2015.02.014

Nandy, A. S., Sharpee, T. O., Reynolds, J. H., & Mitchell, J. F. (2013). The Fine Structure of Shape Tuning in Area V4. Neuron, 78(6), 1102–1115. https://doi.org/10.1016/j.neuron.2013.04.016

Cohen, M. R., & Kohn, A. (2011). Measuring and interpreting neuronal correlations [Number: 7 Publisher: Nature Publishing Group]. Nature Neuroscience, 14(7), 811–819. https://doi.org/10.1038/nn.2842

Power, J. D., Cohen, A. L., Nelson, S. M., Wig, G. S., Barnes, K. A., Church, J. A., Vogel, A. C., Laumann, T. O., Miezin, F. M., Schlaggar, B. L., & Petersen, S. E. (2011). Functional network organization of the human brain. Neuron, 72(4), 665–678. https://doi.org/10.1016/j.neuron.2011.09.006

Haefner, R. M., & Cumming, B. G. (2008). An improved estimator of Variance Explained in the presence of noise. Advances in neural information processing systems, 2008, 585–592.

Adolph, S. C., & Hardin, J. S. (2007). Estimating Phenotypic Correlations: Correcting for Bias Due to Intraindividual Variability [Publisher: [British Ecological Society, Wiley]]. Functional Ecology, 21(1), 178–184.

Sahani, M., & Linden, J. F. (2003). How Linear are Auditory Cortical Responses? In S. Becker, S. Thrun, & K. Obermayer (Eds.). Advances in Neural Information Processing Systems 15 (pp. 125–132). MIT Press.

Pasupathy, A., & Connor, C. E. (2001). Shape Representation in Area V4: Position-Specific Tuning for Boundary Conformation. Journal of Neurophysiology, 86(5), 2505–2519. https://doi.org/10.1152/jn.2001.86.5.2505

Rosner, B., & Willett, W. C. (1988). Interval estimates for correlation coefficients corrected for within-person variation: Implications for study design and hypothesis testing. American Journal of Epidemiology, 127(2), 377–386. https://doi.org/10.1093/oxfordjournals.aje.a114811

Ohzawa, I., Sclar, G., & Freeman, R. D. (1985). Contrast gain control in the cat’s visual system. Journal of Neurophysiology, 54(3), 651–667.

Beaton, G. H., Milner, J., Corey, P., McGuire, V., Cousins, M., Stewart, E., de Ramos, M., Hewitt, D., Grambsch, P. V., Kassim, N., & Little, J. A. (1979). Sources of variance in 24-hour dietary recall data: Implications for nutrition study design and interpretation [Publisher: Oxford Academic]. The American Journal of Clinical Nutrition, 32(12), 2546–2559. https://doi.org/10.1093/ajcn/32.12.2546

Thouless, R. H. (1939). The Effects of Errors of Measurement on Correlation Coefficients [Num Pages: 21 Place: London, etc., United Kingdom, London, etc. Publisher: Cambridge University Press]. British Journal of Psychology. General Section; London, etc., 29(4), 383–403.

Spearman, C. (1904). The proof and measurement of association between two things [Place: US Publisher: Univ of Illinois Press]. The American Journal of Psychology, 15(1), 72–101. https://doi.org/10.2307/1412159

